# Temporal resolution relates to sensory hyperreactivity independently of stimulus detection sensitivity in individuals with autism spectrum disorder

**DOI:** 10.1101/2020.12.23.424156

**Authors:** Ayako Yaguchi, Takeshi Atsumi, Reiko Fukatsu, Masakazu Ide

## Abstract

**Background:** Researchers have been focused on perceptual characteristics of autism spectrum disorder (ASD) in terms of sensory hyperreactivity. Previously, we demonstrated that temporal resolution, which is the accuracy in differentiating the order of two successive vibrotactile stimuli, is associated with the severity of sensory hyperreactivity (Ide et al. 2019). Herein, we examined whether an increase in the perceptual intensity of a tactile stimulus, despite its short duration, is derived from high temporal resolution and a high frequency of sensory temporal summation.

**Method:** Sixteen participants with ASD and fifteen typically developing (TD) participants performed two psychophysical experimental tasks, and we evaluated the detectable duration of vibrotactile stimuli with the same amplitude and temporal resolution. Sensory hyperreactivity was estimated using a self-reported questionnaire.

**Results:** There was no relationship between temporal resolution and the duration of detectable stimuli in either group. However, the ASD group showed more severe sensory hyperreactivity in daily life than the TD group did, and ASD participants with severe sensory hyperreactivity tended to have high temporal resolution but not high sensitivity for detectable duration.

**Conclusion:** Contrary to our hypothesis, there might be different processing between temporal resolution and sensitivity for stimulus detection. Sensory reactivity in daily life would not be based on sensitivity for stimulus detection measured in an experimental task, and we suggest that atypical temporal processing would affect sensory reactivity in ASD. Keywords (within 6 keywords): Temporal resolution, Temporal summation, Autism spectrum disorder, Hyperreactivity, Gap detection

## Introduction

The majority of individuals with autism spectrum disorder (ASD) suffer from atypical sensory experience, which is defined as a diagnostic feature in the Diagnostic and Statistical Manual of Mental Disorders, Fifth Edition (American Psychiatric Association 2013). Sensory reactiveness is understood to be composed of several sub-components, such as over-responsiveness, under-responsiveness, and unique preference for sensory stimuli (Dunn 1997).

Excessively high or low sensitivity to sensory signals has been conventionally assumed to be related to a wide range of individual differences in sensory reactivities (Bryant et al. 2019). For example, individuals with ASD were reported to have lower detection thresholds (or higher sensitivity) for vibrotactile stimuli than that of typically developing (TD) individuals (Blakemore et al. 2006; Cascio et al. 2008). Puts et al. (2014) demonstrated that individuals with ASD have higher detection thresholds, but other studies showed no such difference (Güçlü et al. 2007; Ide et al. 2019). In addition to this inconsistency, some studies reported that the degree of complaints of sensory reactivity estimated using a Sensory Profile, a self-evaluation questionnaire based on Dunn’s theory (Dunn 1997) for adolescents and adults (Adolescent/Adult Sensory Profile: AASP; Brown et al. 2001), was not related to the individual differences in vibrotactile (Ide et al. 2019) or visual detection thresholds (Schulz and Stevenson 2019).

To understand the mechanisms underlying heightened perceptual impressions of stimuli, temporal processing may be an important factor. Both neuronal and perceptual thresholds reduction occur by stimulus inputs over time (Heil and Neubauer 2003). One study tested temporal resolution of visual stimuli by using a simultaneity judgement (SJ) task, where the participants judged whether two successive stimuli divided by a short temporal lag were synchronous or asynchronous (Falter et al. 2012). Individuals with ASD judged two successive stimuli to be asynchronous if they were presented with much shorter temporal lags than TD individuals; that is, better temporal resolution was shown. The high temporal resolution also indicated that they could more precisely judge temporal orders of successive 25-Hz vibrotactile stimuli in a temporal order judgment (TOJ) task compared with that of their TD peers (Tommerdahl et al. 2008). An individual with ASD showed extremely high temporal resolution in a TOJ task of vibrotactile stimuli; the resolution was approximately 10-fold better than that of TD controls, and stronger neural activities in the left ventral premotor cortex (vPMC) were identified as neural substrates of the prominent temporal resolution (Ide et al. 2020). Another study demonstrated that individuals with ASD with high visual temporal resolution in an SJ task tended to have severe social communication deficits and stereotypic behavior as characteristic ASD symptoms, although there was no difference in temporal resolution between ASD and TD groups (Isaksson et al. 2018). In addition, our previous study (Ide et al. 2019) reported no group difference in temporal resolution itself, but individual differences in temporal resolution indexed by TOJ were associated with severity of symptomatic sensory hyperreactivity only in the ASD group: patients with higher temporal resolution of two vibrotactile stimuli complained of severer sensory hyperreactivity scored by the AASP.

Studies of time-based perception imply that high temporal resolution is associated with ASD symptoms, especially severe conditions of sensory hyperreactivity in ASD. However, little detailed explanation has been provided. We proposed that a higher frequency of sensory processing resulting from high firing rates of neurons is associated with a higher temporal resolution of sensory stimuli as atypical synaptic inhibitory functions in the brain have been suggested as a core symptomatic condition of sensory motor dysfunction in ASD (for references of human studies, Foss-Feig et al. 2017; Gaetz et al. 2014; Sapey-Triomphe et al. 2019; Umesawa et al. 2020). The strength of subjective impressions of stimuli would increase with a longer stimulus duration (temporal summation in perception; Barlow 1972), and such threshold reduction may be mediated at the neuronal level (Heil and Neubauer 2003). This leads to a simple prediction that a person with high temporal resolution would perceive a stronger stimulus intensity even over a short stimulus duration, and such association might be explicit in sensory diversion within individuals with ASD.

In the current study, we aimed to answer the following three questions: whether individuals with ASD and high temporal resolution of sensory stimuli can detect tactile stimuli delivered at a near-threshold amplitude, despite a short presentation duration; whether a higher sensitivity to sensory stimuli despite a short presentation duration relates with the severity of sensory hyperreactivity assessed by AASP; and whether our previous finding (Ide et al. 2019) of an association between temporal resolution of vibrotactile stimuli and sensory hyperreactivity is task-independent. For this purpose, we subjected each participant to a systematic test involving two experiments: one to evaluate tactile detectable duration and one to evaluate temporal resolution.

## Methods

Sixteen individuals clinically diagnosed with ASD (mean age, 20.87 years; range, 16–25 years) and fifteen TD individuals (mean age, 19.40 years; range, 16–25 years) were recruited (Table 1). Intelligence quotients (IQs) were estimated using the Wechsler Adult Intelligence Scale-Third Edition (WAIS-III). The Wechsler Intelligence Scale for Children-Fourth Edition was used to evaluate the IQs of one 18-year-old participant (verbal IQ = 88, performance IQ = 89, full scale IQ = 87) whose IQ had been taken three years before the experiment. There was no significant group difference in verbal IQ (*t* (28) = –0.98, *p* = 0.34, Cohen’s *d* = –0.36) or full-scale IQ (*t* (28) = –1.62, *p* = 0.12, Cohen’s *d* = –0.59), while the performance IQ of the ASD group was slightly lower than that of the TD group (*t* (28) = –2.12, *p* = 0.04, Cohen’s *d* = –0.78).

**Table 1.**
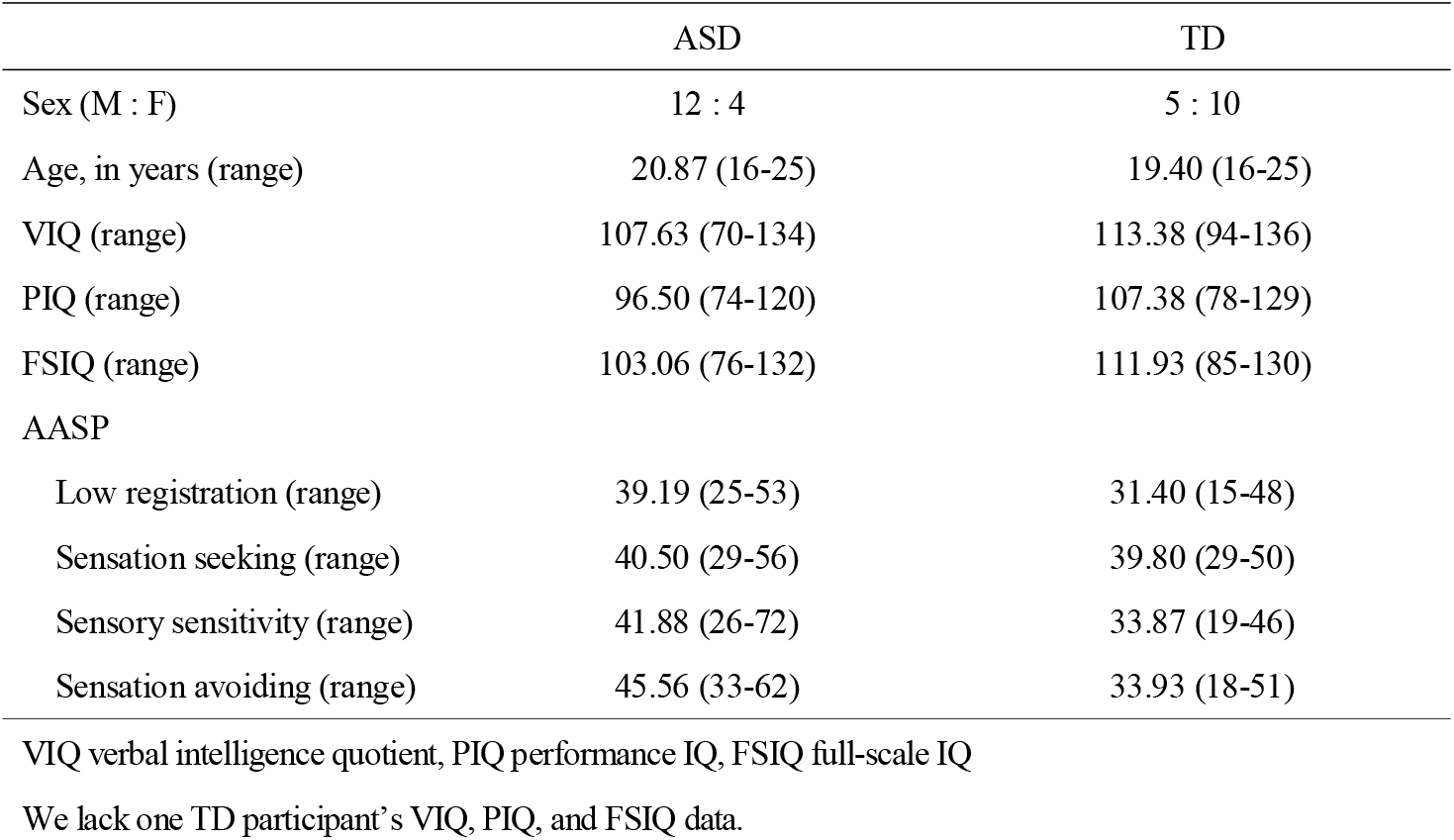
Participant information

|  | ASD | TD |
| --- | --- | --- |
| Sex (M : F) | 12 : 4 | 5 : 10 |
| Age, in years (range) | 20.87 (16-25) | 19.40 (16-25) |
| VIQ (range) | 107.63 (70-134) | 113.38 (94-136) |
| PIQ (range) | 96.50 (74-120) | 107.38 (78-129) |
| FSIQ (range) | 103.06 (76-132) | 111.93 (85-130) |
| AASP |  |  |
| Low registration (range) | 39.19 (25-53) | 31.40 (15-48) |
| Sensation seeking (range) | 40.50 (29-56) | 39.80 (29-50) |
| Sensory sensitivity (range) | 41.88 (26-72) | 33.87 (19-46) |
| Sensation avoiding (range) | 45.56 (33-62) | 33.93 (18-51) |

VIQ verbal intelligence quotient, PIQ performance IQ, FSIQ full-scale IQ
We lack one TD participant's VIQ, PIQ, and FSIQ data.

A Piezo skin contactor (S-16023a, Takei Scientific Instruments; Niigata, Japan) was used to deliver vibrotactile stimulation (50 Hz) on the ventral surface of the left index finger.

### Pre-test: Estimating individual detection threshold of amplitude

We first asked the participants whether they perceived vibrotactile stimuli of varying amplitudes (0, 1, 5, 9, 13, 17, or 21 μm) (Fig. 1: top). Each stimulus condition was repeated 12 times. The response probabilities according to stimulus amplitudes were fitted to a sigmoid function, and the 75% point on the fitting curve was defined as the detection threshold.

**Fig. 1.**
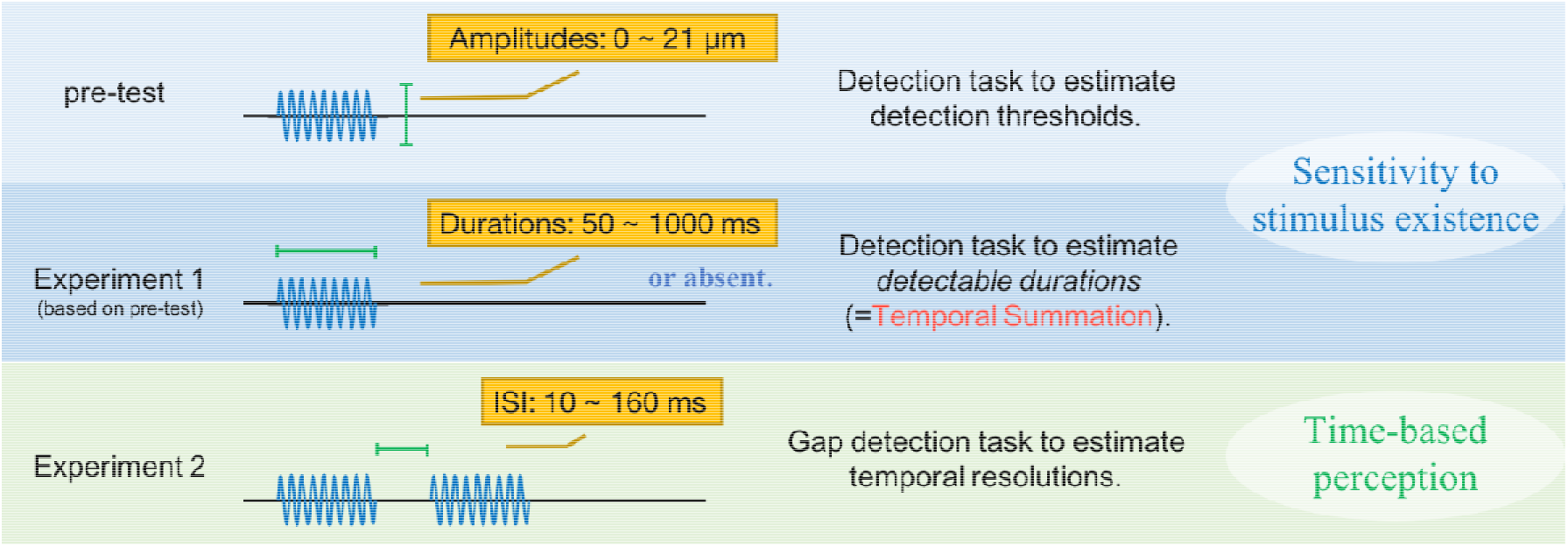
Summary of experimental tasks. (top) Individual detection threshold as the sensitivity to the existence of vibrotactile stimuli was measured. (middle) Detectable vibrotactile stimulus duration was estimated with individually estimated detection thresholds from the pre-test. (bottom) The gap-detection task was introduced to estimate individual temporal resolution of two separate vibrotactile stimuli with varying inter-stimulus intervals (ISIs).

### Experiment 1: Estimating individual detection threshold of duration

We then determined seven durations (50, 100, 200, 400, 600, 800, or 1000 ms) for delivering the stimulus with the defined amplitude for each individual based on the results of the preceding detection task (detection threshold modulated by amplitude) (Fig. 1: middle). The participants were asked whether they perceived stimuli of varying durations with the same stimulus amplitude (i.e., stimulus amplitude was consistently provided at the detection threshold of each participant throughout the test). Each stimulus condition was repeated 12 times. Half of all trials did not deliver any stimulus. The responses associated with each duration were fitted to an exponent function. We defined the 75% point on the fitting curve as the detection threshold along with stimulus duration (hereinafter referred to as the “detectable duration”).

### Experiment 2: Estimating individual temporal resolution

Temporal resolution was estimated using a gap-detection task (Humes et al. 2009) (Fig. 1: bottom). A set of double vibrotactile stimuli (50 Hz) was consecutively delivered to the left index finger at various inter-stimulus intervals (ISI) (10, 40, 70, 100, 130, or 160 ms). Participants were asked about the number of stimuli (1 or 2) detected. Participants’ responses associated with each ISI were fitted to a sigmoid function (Wichmann and Hill 2001) to calculate temporal resolution (75% point).

We evaluated the degree of responsiveness to sensory stimuli using AASP (Brown et al. 2001) that consisted of four sub-categories: low registration, sensation seeking, sensory sensitivity, and sensation avoiding. The former two categories represent lower responsiveness to sensory stimuli, while the latter two categories represent excessive responsiveness.

All procedures performed in studies involving human participants were in accordance with the ethical standards of the ethics committee of the National Rehabilitation Center for Persons with Disabilities and with the 1964 Helsinki Declaration and its later amendments or comparable ethical standards. All participants and their parents gave written informed consent after the study procedures had been fully explained.

## Results

The ASD group showed higher AASP scores than the TD group did for low registration (*F*(1, 27) = 6.57, *p* = 0.01, partial η^2^ = 0.20), sensory sensitivity (*F*(1, 27) = 6.95, *p* = 0.01, partial η^2^ = 0.20), and sensory avoidance (*F*(1, 27) = 14.65, *p* = 0.0002, partial η^2^ = 0.35), but not for sensation seeking (*F*(1, 27) = 0.05, *p* = 0.82, partial η^2^ =0.002) (Fig. 2A). We found no significant difference in the detectable duration or temporal resolution of vibrotactile stimuli between the ASD and TD groups (detectable duration: *t* (29) = 0.76, *p* = 0.45, Cohen’s *d* = 0.11; temporal resolution: *t* (29) = 0.74, *p* = 0.47, Cohen’s *d* = 0.28).

**Fig. 2.**
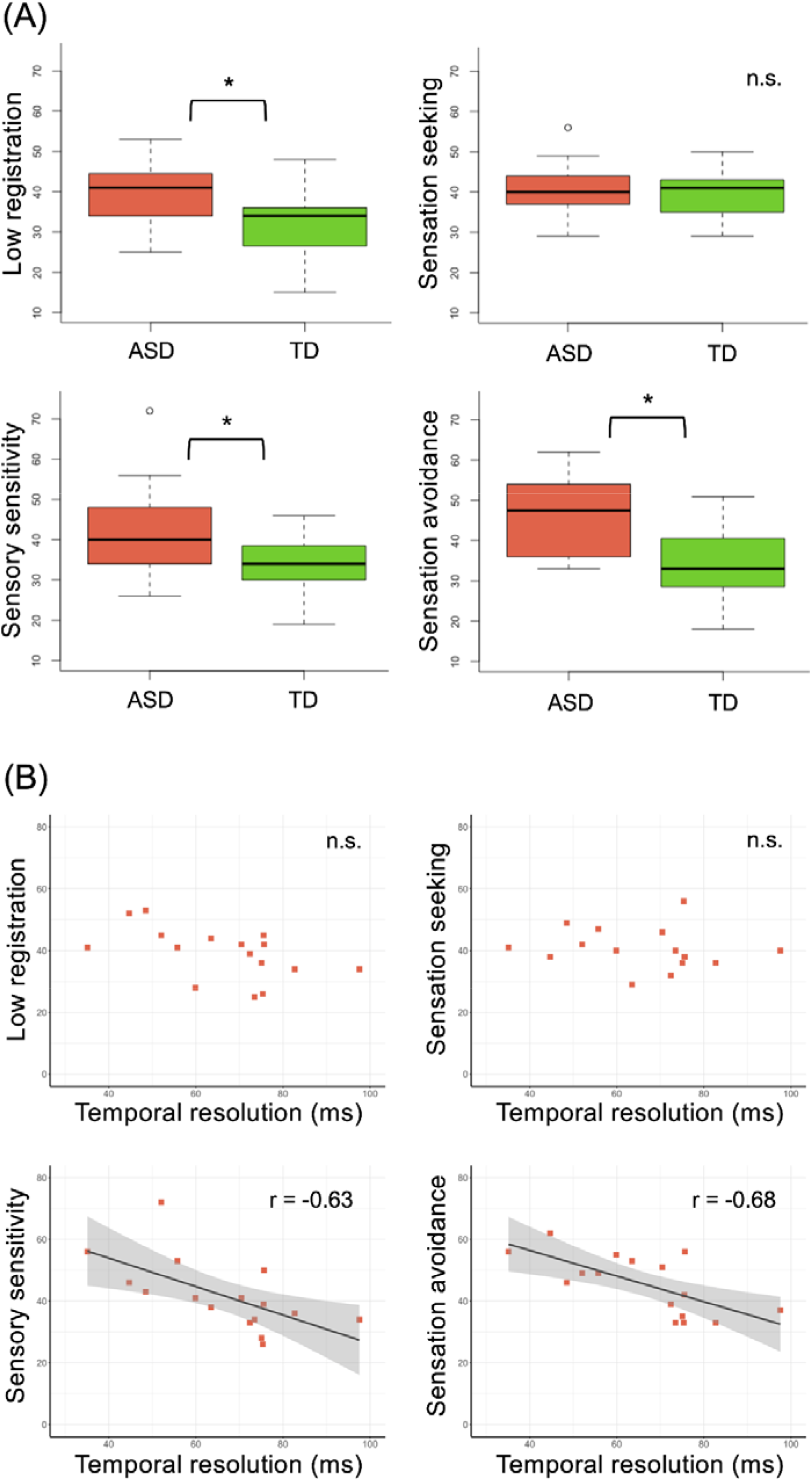
(A) Boxplot of Adolescent/Adult Sensory Profile (AASP) subscale scores. Significant group differences were found in low registration, sensory sensitivity, and sensation avoiding categories (* *p* < 0.05). (B) Scatter plots between temporal resolution and AASP scores in individuals with autism spectrum disorder (ASD). Significant relationships were found in Sensory sensitivity and Sensation avoiding. TD, typically developing.

To confirm whether individual differences in temporal resolution were associated with the detectable duration of stimuli, we performed Pearson’s product-moment correlation coefficient analyses between them. There was no significant correlation in either the ASD (*r* = 0.09, *p* = 0.73, power (1 – β) = 0.06) or TD (*r* = 0.43, *p* = 0.11, power (1 – β) = 0.40) group.

We performed correlation analyses between detectable duration and AASP scores. There was no significant correlation in either the ASD or TD group (ASD group: low registration, *r* = 0.02, *p* = 0.95, power (1 – β) = 0.05; sensation seeking, *r* = –0.27, *p* = 0.32, power (1 – β) = 0.17; sensory sensitivity, *r* = 0.26, *p* = 0.34, power (1 – β) = 0.16; sensation avoidance, *r* = 0.21, *p* = 0.44, power (1 – β) = 0.12 and TD group: low registration, *r* = 0.001, *p* = 0.997, power (1 – β) = 0.05; sensation seeking, *r* = –0.23, *p* = 0.41, power (1 – β) = 0.13; sensory sensitivity, *r* = 0.09, *p* = 0.76, power (1 – β) = 0.06; sensation avoidance, *r* = 0.16, *p* = 0.58, power (1 – β) = 0.09).

We also conducted correlation analyses between temporal resolution and AASP scores (Fig. 2B). Temporal resolution negatively correlated with sensory sensitivity score (*r* = –0.63, *p* = 0.01, power (1 – β) = 0.79) and sensation avoidance score (*r* = –0.68, *p* = 0.003, power (1 – β) = 0.88) in the ASD group but not in the TD group (sensory sensitivity: *r* = 0.13, *p* = 0.65, power (1 – β) = 0.06; sensation avoidance: *r* = –0.04, *p* = 0.90, power (1 – β) = 0.06). This indicates that individuals with ASD and high temporal resolution showed more severe hyperreactivity for daily sensory experience. There was no correlation between temporal resolution and other AASP scores in either the ASD (low registration: *r* = –0.52, *p* = 0.04, power (1 – β) = 0.57, sensation seeking: *r* = –0.15, *p* = 0.58, power (1 – β) = 0.08) or TD group (low registration: *r* = –0.29, *p* = 0.30, power (1 – β) = 0.12, sensation seeking: *r* = –0.43, *p* = 0.11, power (1 – β) = 0.38).

## Discussion

The current study utilized the gap-detection task to evaluate temporal resolution and the stimulus-detection task which manipulated stimulus duration. We investigated whether individuals with ASD and high temporal resolution of sensory stimuli can detect tactile stimuli delivered at a near-threshold amplitude despite a short presentation duration (i.e., detectable duration). We also sought to determine whether there is a relationship between high sensitivity for a short duration stimulus and atypical sensory reactivity. We found no group differences between ASD and TD participants in terms of temporal resolution or detectable duration. Contrary to our expectations, the individual degree of temporal resolution was not related with detectable duration in either group; that is, participants with lower detection thresholds of stimuli at short durations did not exhibit higher temporal resolution of those stimuli. Participants with ASD, but not TD participants, that displayed high temporal resolution in the gap-detection task tended to manifest eminent scores in sensory sensitivity and sensory avoiding categories in AASP, which is congruent with a previous study (Ide et al. 2019), although the scores were not associated with detectable duration in either group.

Based on the idea of temporal summation, duration-sensitive detection thresholds would also explain individual performances in time-perception tasks and sensory hyperreactivity. A previous study reported greater neural activation in the primary somatosensory cortex (SI) and posterior parietal and anterior insular cortices during a vibrotactile detection task than that seen with passive stimulation (Albanese et al. 2009). It has been shown that the intensity of a tactile stimulus is encoded in the SI (Case et al. 2016). However, temporal resolution of a discrimination task of a single or double somatosensory stimulus did not correlate with the peak frequency (the frequency with the highest spectral power) in the SI (Baumgarten et al. 2017). Instead, our previous study demonstrated that task performance in tactile TOJ was associated with neural activation of the right inferior frontal gyrus in both ASD and TD participants (Ide et al. 2020). So far, it has been elucidated that time-based perception demands broader brain functions, such as working memory and interval timing (Buhusi and Meck 2005). Contrary to the tight association of threshold changes in the sensory cortex and stimulus detection (Heil and Neubauer 2003), a simple detection task might not be appropriate to examine whether the perceptual temporal summation framework could adapt to time-based perception, which demands more global neural circuitry.

Sensitivity to sensory stimuli and corresponding reactivity would represent different aspects of sensory processing, so that the reactivity links to how we behaviorally and emotionally react to stimuli after perception (Schulz and Stevenson 2019). Puts and colleagues reported that children with ASD had lower detection thresholds of vibrotactile stimuli than TD children do, although no group difference was found regarding temporal resolution in a TOJ task in the same samples (Puts et al. 2014). In line with this finding, we previously reported that accuracy only in the TOJ task, not in the detection task, is associated with the degree of sensory hyperreactivity (Ide et al. 2019). The previous study showed that detection sensitivity to visual stimuli was not associated with the severity of sensory hyperreactivity measured by Sensory Profile in individuals with ASD but rather with restricted and repetitive behavior (Schulz and Stevenson 2019, Ide et al. 2019). Taking these finding into account, the current results suggested that different underlying mechanisms should be assumed regarding the temporal processing of stimuli and detecting stimuli themselves, especially in relation to sensory hyperreactivity experienced in the daily life of individuals with ASD.

The association between temporal resolution of a flow of multiple stimuli and sensory hyperreactivity was determined to be task-independent because the TOJ and gap-detection tasks were used as the indices in the previous (Ide et al. 2019) and the current study. Further, this suggests that atypical daily sensory experience in ASD is linked to time perception over multiple processing stages. The existence of different stages in terms of human temporal processing in perception/cognition has been described by a two-stage model (Hirsh and Sherrick 1961; Jaśkowski 1991). The first stage is labeled as the “simultaneity center,” which involves generation of judgement on whether sensory stimuli are synchronous or asynchronous, and the second stage (the “order center”) orders sensory stimuli. While the gap-detection task could be assumed to need only the first stage, the TOJ task could require first- and second-stage processing. Indeed, some studies using functional magnetic resonance imaging demonstrated that strong neural responses can be observed in higher-order brain regions including the bilateral frontal and parietal cortices during a TOJ task (Binder 2015; Davis et al. 2009; Takahashi et al. 2013). Moreover, an ASD individual with sensory hyperreactivity who manifested much superior temporal resolution than that of controls in a TOJ task demonstrated greater neural activity in the left vPMC (Ide et al. 2020) where a substantial degree of TOJ-related neural correlates had been reported (Miyazaki et al. 2016). Some studies reported that brain areas, such as the putamen and anterior/posterior insular cortices, which are known to be involved in relatively lower-level sensory processing, are activated when performing SJ, which is hypothesized to relate generally to the first stage of temporal processing (Kimura et al. 2019; Miyazaki et al. 2016). The gap-detection task that we employed in this study would be associated with a relatively earlier stage of temporal processing since the participants simply judged whether the stimuli were perceived as a singlet or a doublet. Given that a previous study (Ide et al. 2019) suggested temporal resolution estimated by the TOJ task is linked to the severity of sensory hyperreactivity in ASD, some components across a broad range of temporal processing might engage with individual degree of the pathological condition of sensory hyperreactivity seen in individuals with ASD.

Enhanced functions of time-based perception and daily sensory experiences in ASD might not be induced by a greater amplitude of neural activation in specific brain regions but rather by the atypical functional connectivity mediated by rhythmic neural activations within the relating circuits. Recent advances in neuroscientific research have been demonstrated that perceptual processing cycle is modulated by phasic neural oscillation, engaging large individual differences of stimulus temporal resolution. The phase of an alpha- or low beta-oscillation (8–20 Hz) in SI predicted discrimination accuracy of electrical pulse stimulations (Baumgarten et al. 2015). There is another report that the occipital alpha frequency recorded by electroencephalography shortly before the stimulus presentation can predict temporal resolution of perceived frequency of flickering visual stimuli (Samaha and Postle 2015). As assuming a phasic neural oscillation reflects cycle of perceptual processing, stimulus temporal resolution would be modulated by the cycle. The study of tactile TOJ with using magnetoencephalography (MEG) demonstrated that the phase of alpha-band oscillation is crucial for proper performances of TOJ (Takahashi and Kitazawa 2017). One research suggested that the left vPMC is the important neural correlates of TOJ (Miyazaki et al. 2016). Our previous study using the proton magnetic resonance spectroscopy (^1^H-MRS) found that individuals with ASD who showed reduced concentration of an inhibitory metabolite, γ-aminobutyric acid (GABA) in the left vPMC tended to argue severer sensory hyperreactivity estimated by AASP (Umesawa et al. 2020). Considering these findings, we assume that decreased neural inhibition by GABA-mediated atypical neural function, which has been broadly reported across ASD studies (see, Cellot and Cherubini 2014; Pizzarelli and Cherubini 2011), in the brain circuits of temporal resolution might associate with extraordinarily high perceptual processing cycle and result in frequent experience of sensory hyperreactivity. GABA plays an important role for generating neural oscillation. Rhythmic inhibitory postsynaptic potential (IPSP) provided by GABAergic interneurons, such as parvalbumin (PV) neuron could synchronize neuronal spikes, and this leads to an entraining of neural oscillation (Cobb et al. 1995; Muthukumaraswamy et al. 2015; Wang and Buzsáki 1996). We speculate that GABAergic dysfunction alters neural oscillations across brain circuits associated with time-based perception and sensory reactivity, inducing each enhanced function.

We concluded that temporal resolution measured by several methods reflecting different stages of temporal processing of stimuli might be associated with individual degrees of sensory hyperreactivity. Temporal resolution in the task like we employed could be modulated by neural oscillation rather than the simple amplitude of neural responses, but it is still unknown whether time-based perception directly amplifies the perceptual intensity of a stimulus in ASD. One possibility is that phasic neural oscillation, which modulates temporal resolution of successive stimuli, also alters perceptual intensity by inducing a summation of the perceptual cycle rather than reducing the threshold. Further studies are needed to understand the dissociative mechanism between time-based perception and detection sensitivity modulated by temporal properties of sensory stimuli that contributes to pathological states of sensory hyperreactivity in ASD.

## Acknowledgements

This study was supported by Japan Society for the Promotion of Science (No.JP18J22022, No.JP18H03663, No.JP18H03140, No.JP17942790). We would like to thank Yumi Umesawa and Yichin Wang for their technical help.

## References

Albanese, M.-C., Duerden, E. G., Bohotin, V., Rainville, P., & Duncan, G. H. (2009). Differential Effects of Cognitive Demand on Human Cortical Activation Associated With Vibrotactile Stimulation. Journal of Neurophysiology, 102(3), 1623–1631.

American Psychiatric Association (2013). Diagnostic and Statistical Manual of Mental Disorders (DSM-5). American Psychiatric Press, Inc., Washington, DC, USA.

Barlow, H. B. (1972). Single units and sensation: a neuron doctrine for perceptual psychology? Perception, 1(4), 371–394.

Baumgarten, T. J., Schnitzler, A., & Lange, J. (2015). Beta oscillations define discrete perceptual cycles in the somatosensory domain. Proceedings of the National Academy of Sciences, 112(39), 12187–12192.

Baumgarten, T. J., Schnitzler, A., & Lange, J. (2017). Beyond the Peak – Tactile Temporal Discrimination Does Not Correlate with Individual Peak Frequencies in Somatosensory Cortex. Frontiers in Psychology, 8.

Binder, M. (2015). Neural correlates of audiovisual temporal processing–comparison of temporal order and simultaneity judgments. Neuroscience, 300, 432–447.

Blakemore, S. J., Tavassoli, T., Calo, S., Thomas, R. M., Catmur, C., Frith, U., et al. (2006). Tactile sensitivity in Asperger syndrome. Brain and cognition, 61(1), 5–13, doi:10.1016/j.bandc.2005.12.013.

Brown, C., Tollefson, N., Dunn, W., Cromwell, R., & Filion, D. (2001). The adult sensory profile: Measuring patterns of sensory processing. The American Journal of Occupational Therapy, 55(1), 75–82.

Bryant, L. K., Woynaroski, T. G., Wallace, M. T., & Cascio, C. J. (2019). Self-reported Sensory Hypersensitivity Moderates Association Between Tactile Psychophysical Performance and Autism-Related Traits in Neurotypical Adults. Journal of autism developmental disorders, 1―14.

Buhusi, C. V., & Meck, W. H. (2005). What makes us tick? Functional and neural mechanisms of interval timing. Nature Reviews Neuroscience, 6(10), 755–765.

Cascio, C. J., McGlone, F., Folger, S., Tannan, V., Baranek, G., Pelphrey, K. A., et al. (2008). Tactile perception in adults with autism: a multidimensional psychophysical study. Journal of Autism and Developmental Disorders, 38(1), 127–137.

Case, L. K., Laubacher, C. M., Olausson, H., Wang, B., Spagnolo, P. A., & Bushnell, M. C. (2016). Encoding of Touch Intensity But Not Pleasantness in Human Primary Somatosensory Cortex. Journal of Neuroscience, 36(21), 5850–5860.

Cellot, G., & Cherubini, E. (2014). GABAergic signaling as therapeutic target for autism spectrum disorders. Frontiers in pediatrics, 2, 70.

Cobb, S. R., Buhl, E. H., Halasy, K., Paulsen, O., & Somogyi, P. (1995). Synchronization of neuronal activity in hippocampus by individual GABAergic interneurons. Nature, 378(6552), 75–78.

Davis, B., Christie, J., & Rorden, C. (2009). Temporal order judgments activate temporal parietal junction. Journal of Neuroscience, 29(10), 3182–3188.

Dunn, W. (1997). The impact of sensory processing abilities on the daily lives of young children and their families: A conceptual model. Infants and young children, 9, 23–35.

Falter, C. M., Elliott, M. A., & Bailey, A. J. (2012). Enhanced visual temporal resolution in autism spectrum disorders. PLoS One, 7(3).

Güçlü, B., Tanidir, C., Mukaddes, N. M., & Ünal, F. (2007). Tactile sensitivity of normal and autistic children. Somatosensory & motor research, 24(1―2), 21–33.

Heil, P., & Neubauer, H. (2003). A unifying basis of auditory thresholds based on temporal summation. Proceedings of the National Academy of Sciences of the United States of America, 100(10), 6151–6156.

Hirsh, I. J., & Sherrick Jr, C. E. (1961). Perceived order in different sense modalities. Journal of experimental psychology, 62(5), 423.

Humes, L. E., Busey, T. A., Craig, J. C., & Kewley-Port, D. (2009). The effects of age on sensory thresholds and temporal gap detection in hearing, vision, and touch. Attention, Perception, Psychophysics, 71(4), 860–871.

Hirsh, I. J., & Sherrick, C. E., Jr. (1961). Perceived order in different sensory modality. Journal of Experimental Psychology, 62(5), 423–432.

Ide, M., Atsumi, T., Chakrabarty, M., Yaguchi, A., Umesawa, Y., Fukatsu, R., & Wada, M. (2020). Neural basis of extremely high temporal sensitivity: Insights from a patient with autism. Frontiers in Neuroscience, 14 340.

Ide, M., Yaguchi, A., Sano, M., Fukatsu, R., & Wada, M. (2019). Higher tactile temporal resolution as a basis of hypersensitivity in individuals with autism spectrum disorder. Journal of Autism and Developmental Disorders, 49(1), 44–53.

Isaksson, S., Salomäki, S., Tuominen, J., Arstila, V., Falter-Wagner, C. M., & Noreika, V. (2018). Is there a generalized timing impairment in Autism Spectrum Disorders across time scales and paradigms? Journal of psychiatric research, 99 111–121.

Jaśkowski, P. (1991). Two-stage model for order discrimination. Perception & psychophysics, 50(1), 76–82.

Kimura, T., Kadota, H., Kuroda, T., Funai, T. D., Iwata, M., Kochiyama, T., & Miyazaki, M. (2019). Neural correlates of tactile simultaneity judgement: a functional magnetic resonance imaging study. Scientific reports, 9(1), 1–13.

Lederman, S. J., & Klatzky, R. L. (2009). Haptic perception: A tutorial. Attention, Perception, & Psychophysics, 71(7), 1439–1459.

McIntosh, D. N., Miller, L. J., & Shyu, V. (1999). Development and validation of the Short Sensory Profile. In W. Dunn (Ed.), The Sensory Profile examiner’s manual (pp. 59―73). San Antonio, TX: Psychological Corporation.

Muthukumaraswamy, S. D., Shaw, A. D., Jackson, L. E., Hall, J., Moran, R., & Saxena, N. (2015). Evidence that Subanesthetic Doses of Ketamine Cause Sustained Disruptions of NMDA and AMPA-Mediated Frontoparietal Connectivity in Humans. The Journal of Neuroscience, 35(33), 11694–11706.

Pizzarelli, R., & Cherubini, E. (2011). Alterations of GABAergic signaling in autism spectrum disorders. Neural plasticity, 2011.

Puts, N. A., Wodka, E. L., Tommerdahl, M., Mostofsky, S. H., & Edden, R. A. (2014). Impaired tactile processing in children with autism spectrum disorder. Journal of neurophysiology, 111(9), 1803–1811.

Samaha, J., & Postle, B. R. (2015). The speed of alpha-band oscillations predicts the temporal resolution of visual perception. Current Biology, 25(22), 2985–2990.

Schulz, S. E., & Stevenson, R. A. (2019). Differentiating between sensory sensitivity and sensory reactivity in relation to restricted interests and repetitive behaviours. Autism, 1362361319850402.

Takahashi, T., Kansaku, K., Wada, M., Shibuya, S., & Kitazawa, S. (2013). Neural correlates of tactile temporal-order judgment in humans: an fMRI study. Cereb Cortex, 23(8), 1952–1964, doi:10.1093/cercor/bhs179.

Tomchek, S. D., & Dunn, W. (2007). Sensory processing in children with and without autism: a comparative study using the short sensory profile. American Journal of occupational therapy, 61(2), 190–200.

Umesawa, Y., Atsumi, T., Chakrabarty, M., Fukatsu, R., & Ide, M. (2020). GABA concentration in the left ventral premotor cortex associates with sensory hyper-responsiveness in autism spectrum disorders without intellectual disability. Frontiers in Neuroscience, 14 482.

Wang, X.-J., & Buzsáki, G. (1996). Gamma Oscillation by Synaptic Inhibition in a Hippocampal Interneuronal Network Model. The Journal of Neuroscience, 16(20), 6402–6413.

Wichmann, F. A., & Hill, N. J. (2001). The psychometric function: I. Fitting, sampling, and goodness of fit. Perception & psychophysics, 63(8), 1293–1313.

